# Functional diversity of Arabidopsis late embryogenesis abundant proteins in response to changes in the physicochemical environment

**DOI:** 10.1101/2025.08.26.672499

**Authors:** LD Palomino-Navarrete, D Pérez-Villanueva, D Moses, E Gollub, K Nguyen, S Sanchez-Martinez, F Yu, K Martinez, TC Boothby, S Sukenik, CL Cuevas-Velazquez

## Abstract

Some desiccation-tolerant organisms accumulate intrinsically disordered proteins (IDPs), such as Late Embryogenesis Abundant (LEA) proteins, which help protect other proteins from inactivation and/or aggregation during desiccation. Like other IDPs, LEA proteins adopt ensembles of extended conformations that shift in response to environmental changes. Desiccation causes dramatic changes in the cellular environment, but it is unclear how the structural malleability of LEAs is related to their protective function. In this work, we measured the *in vitro* protective function and structural sensitivity to changes in the environment of four *Arabidopsis thaliana* LEA proteins from different families. We found that all LEAs showed different protection efficiencies of the labile enzyme lactate dehydrogenase under desiccation *in vitro*. In line with this, we identified distinct ensemble structural changes when these LEA proteins were exposed to different physicochemical environments. Specifically, AtEM1, AtLEA7, and AtLEA4-5 showed compaction when the solution was crowded with polymers, whereas AtLEA6-2.2 showed larger structural changes when salt concentrations were increased. Furthermore, the ensembles of AtEM1, AtLEA7, and AtLEA4-5 gained helicity under desiccated conditions, while that of AtLEA6-2.2 remained largely disordered. Our results highlight how ensemble properties of LEA proteins contribute to their distinct functional activities *in vitro*. This work advances our understanding of the molecular features that contribute to functional diversity in desiccation-related IDPs.

## 1. INTRODUCTION

Water is essential for all living organisms. Beyond its fundamental role as the universal solvent, it plays a pivotal part in the growth, development, and physiology of land plants (Choat et al., 2018). Due to their stationary nature, plants cannot avoid environmental changes the way that animals do, and must instead develop adaptive mechanisms to cope with them. Water deficit is one of the most significant environmental stressors that plants encounter throughout their life cycle (Bray, 1997). Plants have evolved various strategies to cope with low water availability. One widely conserved response across all plants is the accumulation of a group of proteins known as Late Embryogenesis Abundant (LEA) proteins (Battaglia et al., 2008). LEA proteins accumulate during the last stage of embryogenesis in seed-bearing plants, coinciding with the acquisition of desiccation tolerance. In addition to their role in seeds, LEA proteins are also induced in vegetative tissues under water deficit conditions (Olvera-Carrillo et al., 2010). LEA proteins are known to be key players in tolerance of dehydration and desiccation since mutants lacking some genes that encode for LEA proteins are sensitive to these conditions (Manfre et al., 2006; Olvera-Carrillo et al., 2010; Senthil-Kumar & Udayakumar, 2006). The association between LEA proteins and water-deficit tolerance extends beyond plants, as desiccation-tolerant organisms such as bacteria, nematodes, rotifers, tardigrades, springtails, and insects also possess genes encoding LEA-Like proteins (Hernández-Sánchez et al., 2022).

Despite the crucial role of LEA proteins in plant responses to water limitation, their precise molecular functions and action mechanisms *in vivo* remain poorly understood. *In vitro* assays using model ions, membranes, carbohydrates, nucleic acids, and enzymes support divergent hypotheses, suggesting that LEA proteins may function as radical scavengers, membrane stabilizers, sugar and nucleic acid-binding proteins, and protein protectants (Hundertmark et al., 2011; Liu et al., 2011; Reyes et al., 2005, 2008). The function of LEA proteins as protein stabilizers has been extensively studied over the past decades (Artur et al., 2019; Cuevas-Velazquez et al., 2016; Hernández-Sánchez et al., 2024; Janis et al., 2022; Kc et al., 2024; Lin & Thomashow, 1992; Rendón-Luna et al., 2024; Reyes et al., 2005). LEA proteins prevent the inactivation (and in some cases, the aggregation) of labile enzymes subjected to desiccation *in vitro* (Kovacs et al., 2008; Reyes et al., 2005, 2008).

Most LEA proteins are relatively small (under 200 residues) and are enriched in polar and charged amino acids (Battaglia et al., 2008; Hernández-Sánchez et al., 2022). Because of these properties, LEA proteins are predicted, and in many cases experimentally demonstrated, to be intrinsically disordered proteins (IDPs) (Cuevas-Velazquez et al., 2016; Hundertmark et al., 2012; Popova et al., 2011; Rivera-Najera et al., 2014; Shih et al., 2010). Unlike well-folded proteins, IDPs do not possess a single stable native structure (Wright & Dyson, 2015). Instead, they exist as a dynamic ensemble of interchanging conformations (Holehouse & Kragelund, 2023). Due to their extended and flexible nature, IDPs are particularly sensitive to changes in the physicochemical properties of the solution environment (Moses et al., 2023). This is because in the absence of an extensive network of intramolecular bonds, changes to solvent:IDP interactions can reweigh the preferred structures in the ensemble. For example, when the termini of an IDP containing oppositely charged residues encounters a solution with high ionic strength, masking charges will weaken these interactions, causing the ensemble to expand.

The conformational ensemble of disordered LEA proteins is particularly sensitive to environmental changes, suggesting that they might function as sensors of dehydration/desiccation conditions (Artur et al., 2019; Bremer et al., 2017; Cuevas-Velazquez et al., 2016, 2021; Hernández-Sánchez et al., 2024; Moses et al., 2023; Rivera-Najera et al., 2014). Under conditions of high osmolarity and/or high macromolecular crowding in solution, some LEA proteins undergo a disorder-to-folded transition, specifically adopting helical conformations (Bremer et al., 2017; Cuevas-Velazquez et al., 2016, 2021; Rendón-Luna et al., 2024). A gain in helicity also occurs when some LEA proteins are fully desiccated (Kc et al., 2024; Thalhammer et al., 2015). This conformational sensitivity is thought to underlie their *in vitro* function as protein stabilizers during water deficit stress (desiccation and freeze-thaw cycles) (Artur et al., 2019; Cuevas-Velazquez et al., 2016). For example, AtLEA4-5 from the model plant *Arabidopsis thaliana* (Arabidopsis) possesses an N-terminal conserved domain that can adopt a helical conformation and is necessary and sufficient to protect the labile enzyme lactate dehydrogenase (LDH) from desiccation and freeze-thaw cycles (Cuevas-Velazquez et al., 2016). The addition of proline residues across this domain disrupts the acquisition of helical conformation and reduces its protective efficiency over LDH, but only when the domain is tested in the absence of the C-terminal variable domain (Rendón-Luna et al., 2024). Similar structure-function relationships have been observed in other disordered protectants such as the tardigrade specific cytosolic abundant heat soluble (CAHS) proteins, which are critical for tardigrade (water bears) desiccation tolerance (Boothby et al., 2017). Mutations that disrupted the helical conformation of the linker region in CAHS D from *Hypsibius exemplaris* impaired its ability to protect LDH during desiccation, although it was not the only property responsible for this effect (Biswas et al., 2024).

Most studies on LEA proteins focus on individual proteins from specific families, with only a few systematically comparing the ensemble-function relationship across different LEA groups of the same organism (Artur et al., 2019). Furthermore, variations in experimental methods used to characterize LEA protein’s protective function *in vitro* (i.e., desiccation vs freeze-thaw cycles), make it difficult to directly compare their protective efficiencies across studies.

In this work, we measured the protein stabilization efficiency during desiccation and the ensemble sensitivity to environmental perturbations of four Arabidopsis LEA proteins from group 1 (AtEM1, At3g51810), group 3 (AtLEA7, At1g52690), group 4 (AtLEA4-5, At5g06760), and group 6 (AtLEA6-2.2, At2g23120). We found that these proteins exhibit different protective efficiencies for LDH during desiccation *in vitro* and their conformational ensembles display varying degrees of sensitivity to changes in the surrounding chemical environment and to desiccation. Taken together, our findings suggest that the sequence-encoded molecular features of Arabidopsis LEA protein ensembles contribute to their functional diversity in response to changes in the physicochemical environment *in vitro*, a possibility that could expand to their functions in cells.

## 2. RESULTS

### 2.1 Selection of Arabidopsis LEA proteins from different groups

The Arabidopsis genome encodes 51 LEA proteins from six different groups, which include both hydrophobic (atypical) and hydrophilic (typical) proteins (Hundertmark & Hincha, 2008). To compare the protective and sensing functions of selected Arabidopsis LEA proteins from different groups, we focused on representative proteins whose transcripts are highly abundant and strongly overexpressed in dry seeds (**Figure 1a**). Additionally, we prioritize hydrophilic LEA proteins - those enriched in polar, charged, and small residues, since such IDPs are associated with the protection of labile enzymes during water deficit conditions (Garay-Arroyo et al., 2000; Reyes et al., 2005). Finally, due to the method we used to experimentally quantify the global dimensions of conformational ensembles under different conditions, we selected proteins with less than 170 amino acids (Moses et al., 2020). Based on this criteria, we initially selected five LEA proteins, but only four could be successfully expressed and purified for further characterization: AtEM1 (group 1), AtLEA7 (group 3), AtLEA4-5 (group 4), and AtLEA6-(group 6) (**Figure 1b**).

**FIGURE 1.**
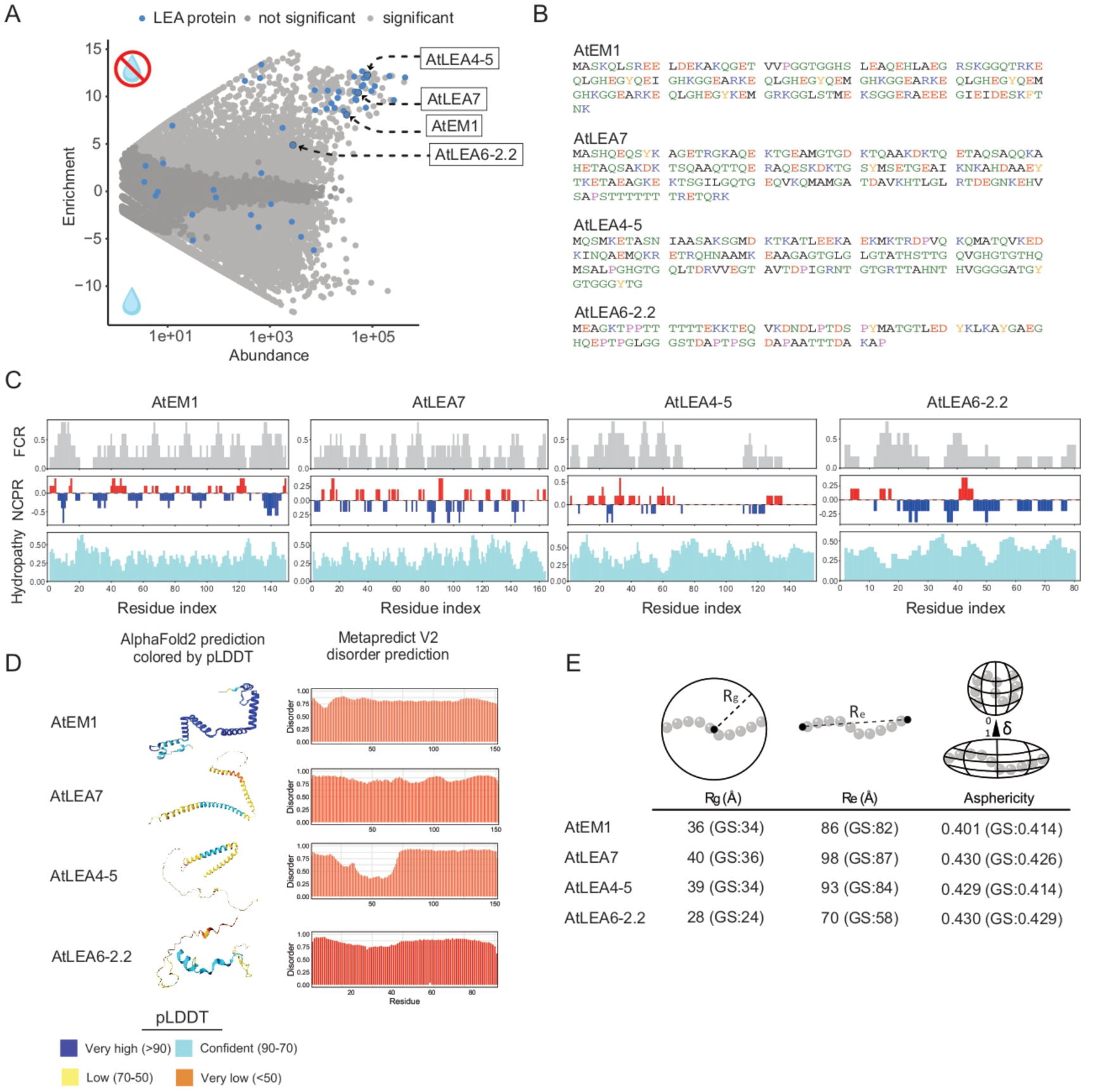
Selected representatives of Arabidopsis LEA proteins are predicted to be disordered with different propensities to form α-helices. A) Differential gene expression analysis of Arabidopsis seeds before and after desiccation. Light gray dots: significant differentially expressed transcripts (*p≤*0.05). Dark gray dots: not significant differentially expressed transcripts (*p*>0.05). Blue dots: transcripts encoding LEA proteins. RNA-seq data obtained from (Klepikova et al., 2016). B) Primary amino acid sequences of AtEM1, AtLEA7, AtLEA4-5, and AtLEA6-2.2. C) Fraction of charged residues (FCR), net charge per residue (NCPR), and hydropathy levels across the sequences of AtEM1, AtLEA7, AtLEA4-5, and AtLEA6-2.2. D) AlphaFold2 predicted tridimensional structure (left) and Metapredict disordered prediction (right) of AtEM1, AtLEA7, AtLEA4-5, and AtLEA6-2.2. E) Average ensemble dimensions of AtEM1, AtLEA7, AtLEA4-5, and AtLEA6-2.2 predicted from ALBATROSS deep-learning model. Predictions of the same parameters for GS repeats of the same lengths are indicated in parenthesis.

The four selected LEA proteins have a fraction of charged residues (FCR) greater than 0.2, an average hydropathy less than 4, a net charge per residue (NCPR) near neutrality, and well-mixed oppositely charged amino acids (*κ* < 0.15) (Holehouse et al., 2015) (**Figure 1c, Table S1**). All four proteins are predicted to be largely disordered throughout their sequence; however, AlphaFold2 models predicted long helices for AtEM1, AtLEA7, and AtLEA4-5 (**Figure 1d**). The predicted properties and conformational preferences of the selected LEA proteins are similar to other desiccation protective proteins such as hydrophilins, tardigrade disordered proteins, and heat-resistant obscure proteins (Biswas et al., 2024; Garay-Arroyo et al., 2000). To further assess the conformational properties of the selected LEA proteins, we predicted their average ensemble dimensions using the deep-learning model ALBATROSS (Lotthammer et al., 2024). We found that the predicted radius of gyration (*R*_g_) ranged from 28 Å to 40 Å, while the predicted end-to-end distance (*R*_e_) ranged from 70 Å to 98 Å (**Figure 1e**). Additionally, all four LEA proteins showed similar predicted asphericity values (**Figure 1e**). In summary, we selected a set of four Arabidopsis LEA proteins from different families, with similar sequence chemistry properties but predicted differences regarding their potential ensemble properties, to evaluate their ensemble-function relationship.

### 2.2 Arabidopsis LEA proteins exhibit different LDH protection efficiencies upon desiccation in vitro

To test the protective function of the selected LEA proteins, we purified recombinant LEA proteins (**Figure S1**), and assessed their ability to preserve lactate dehydrogenase (LDH) activity after desiccation (Reyes et al., 2005). LDH was either completely desiccated without additives or mixed with increasing amounts of each LEA protein before desiccation. After rehydration, we measured the remaining LDH activity. We found that all LEA proteins preserved LDH activity from desiccation-induced inactivation in a concentration-dependent manner (**Figure 2a**). To quantitatively compare their protective efficiencies, we calculated the protective dose 50% (PD50) - the amount of LEA protein required to retain 50% LDH activity after desiccation. We found that all proteins differentially protected LDH from desiccation: AtLEA4-5 was the best protectant, followed by AtEM1, AtLEA6-2.2, and AtLEA7 (**Figure 2b**). Our results indicate that, despite their similar sequence chemistry, the four LEA proteins tested have different protective activities, suggesting that their potential differences in ensemble features might contribute to their protective function *in vitro*.

**FIGURE 2.**
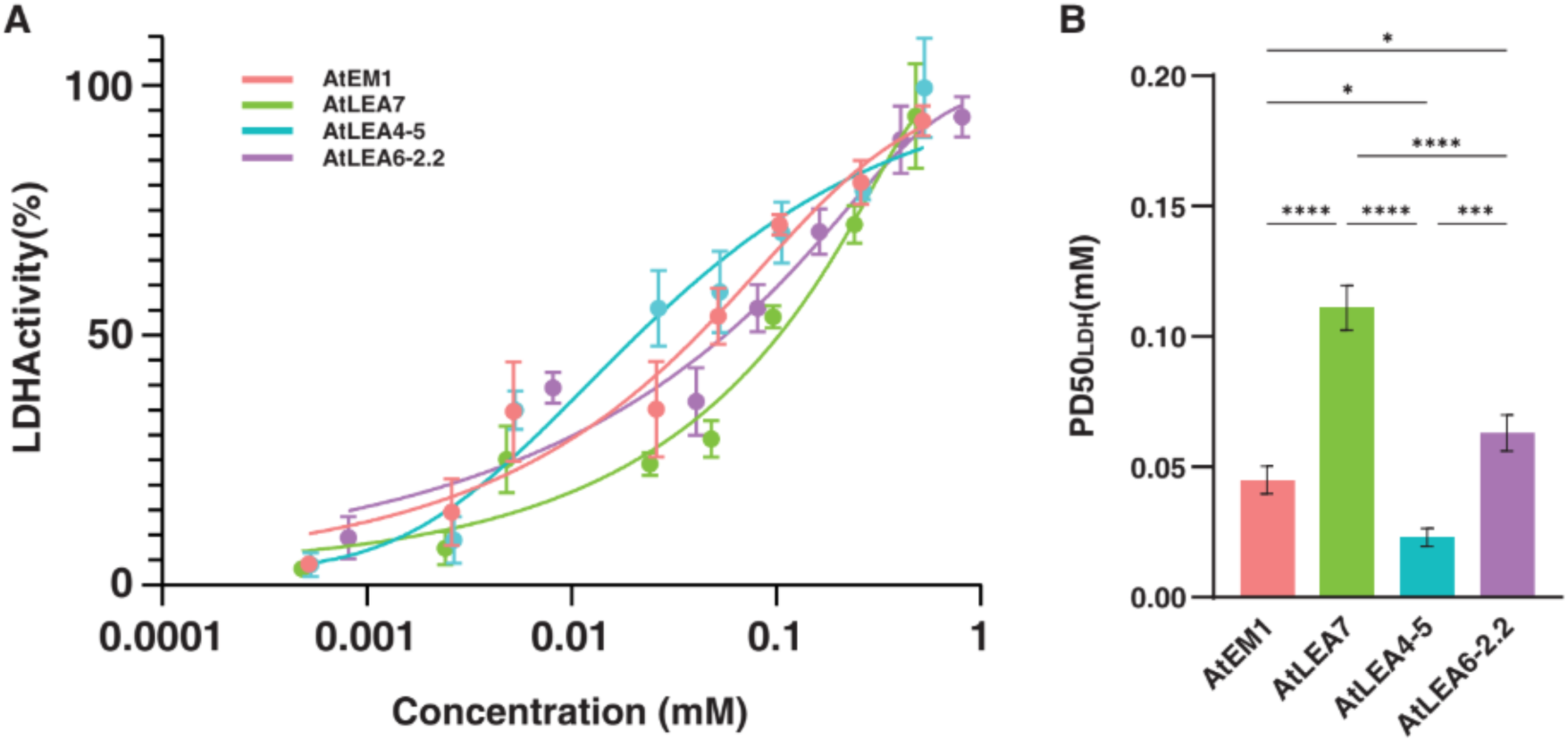
Arabidopsis LEA proteins differentially protect LDH from desiccation *in vitro*. A) Concentration dependence of LDH activity protection by LEA proteins, relative to the untreated sample (100%). Mean ± SD from *n* = 3 independent measurements. B) Protective dose 50% (PD50) of each LEA protein. Mean ± SD from *n* = 3 independent measurements. One-way ANOVA with Tukey’s HSD test. \**p*<0.05, \*\**p*<0.01, \*\*\**p*<0.001, \*\*\*\**p*<0.0001, ns *p*>0.05.

### 2.3 The conformational ensembles of Arabidopsis LEA proteins differ in their sensitivity to solution chemistry

Because of the small number of intramolecular bonds and high degree of surface exposure, some IDPs display structural sensitivity to changes in their solution environment (Moses et al., 2023). To assess the ensemble structural sensitivity of the four LEA proteins to their solution environment, we first characterized their ensemble dimensions *in silico* using all-atom Monte Carlo simulations (Vitalis & Pappu, 2009). Briefly, each sequence is placed in an implicit solvent with increasing repulsive interactions with the backbone. Sequences that are more sensitive to environmental changes will show greater reductions in ensemble dimensions, while more rigid, less sensitive sequences will undergo smaller changes (Holehouse & Sukenik, 2020). In addition, sensitive sequences will show these changes in less repulsive solutions compared to less sensitive ones.

To ensure proper conformational sampling for AtEM1 and AtLEA7, we divided their sequences into two segments with a 20 amino acid overlap between segments. We found that regardless of the segment simulated, the overall sensitivity is retained (**Figure 3**). Thus, AtEM1_1 and AtEM1_2, which comprise the N- and C-terminals of the same protein, respectively, showed similar sensitivity. The same was true for AtLEA7_1 and AtLEA7_2 (**Figure 3**). Among all tested sequences, AtLEA4-5 showed both the highest degree of compaction and the greatest sensitivity to repulsive solutions, as shown by the magnitude of the normalized R*_e_* change and the midpoint of this change on the x-axis, respectively (**Figure 3a**). AtEM1 and AtLEA6-2.2 showed the lowest degree of change and the lowest degree of sensitivity (**Figure 3**).

**FIGURE 3.**
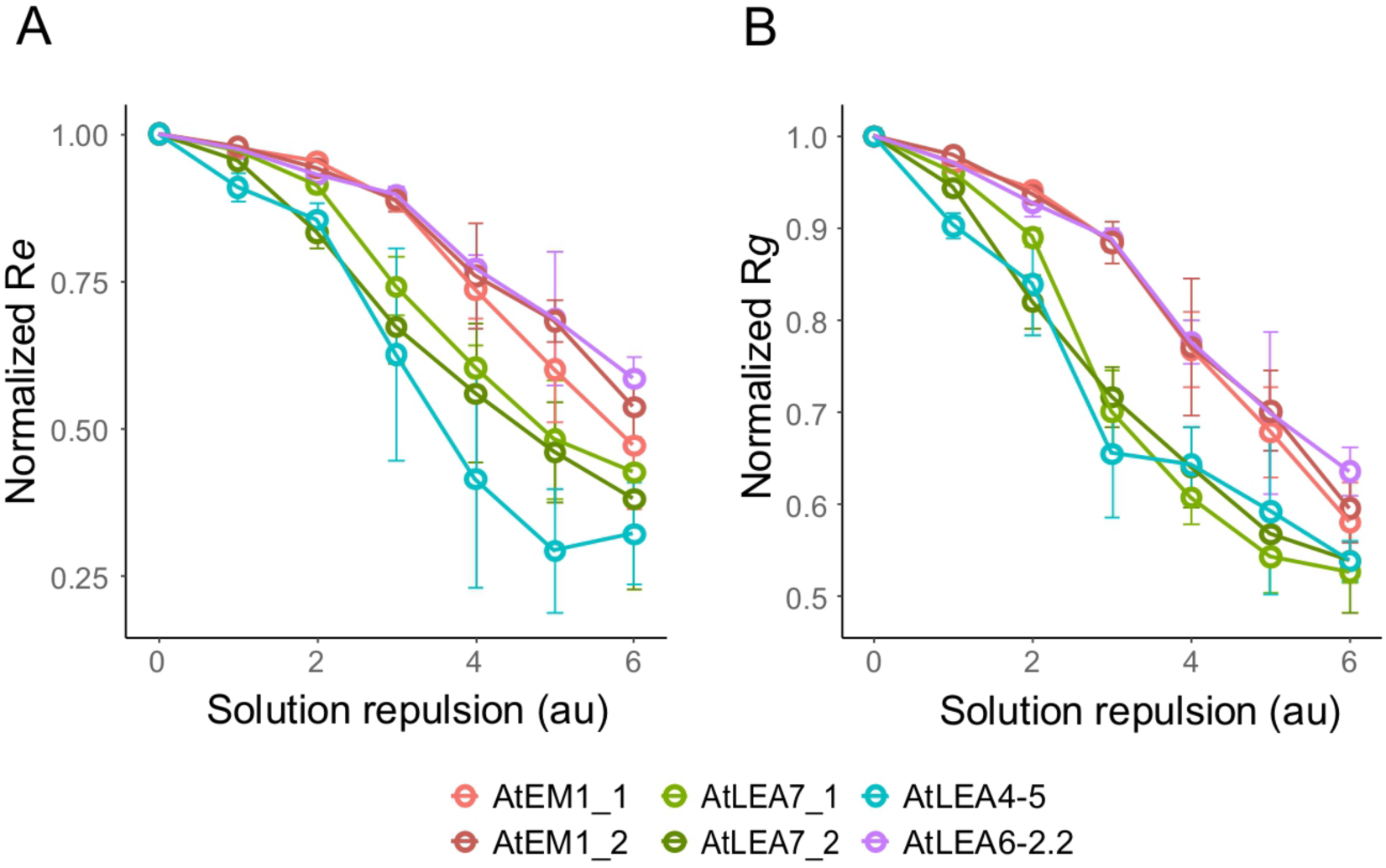
Arabidopsis LEA proteins are highly sensitive to changes in the solution environment *in silico*. All-atom Monte Carlo simulations of selected Arabidopsis LEA proteins. Long sequences (AtEM1 and AtLEA7) were divided into two shorter segments (AtEM1_1, AtEM1_2, AtLEA7_1, AtLEA7_2) with a 20 amino acid overlap between segments. A) Normalized end-to-end distance (*R_e_*) of AtEM1_1, AtEM1_2, AtLEA7_1, AtLEA7_2, AtLEA4-5, and AtLEA6-2.2 in simulated solutions with increasing repulsion to the protein backbone. B) Normalized radii of gyration (*R_g_*) of AtEM1_1, AtEM1_2, AtLEA7_1, AtLEA7_2, AtLEA4-5, and AtLEA6-2.2 in simulated solutions with increasing repulsion to the protein backbone. Each normalized *R_e_* and *R_g_* was normalized based on the most expanding solution (solution repulsion = 0 arbitrary units) to highlight solution sensitivity. Mean ± SD from *n* = 5 independent simulations.

We next wanted to determine if these trends hold *in vitro*. To do so, we performed solution space scanning (SSS), a method that has previously been used to track the structural sensitivity of IDP ensembles across different chemical environments (Cuevas-Velazquez et al., 2021; Moses et al., 2020, 2024). For this, an IDP of interest is placed between a donor (mTurqouise2) and an acceptor fluorescent protein (mNeonGreen). These fused proteins form a Förster resonance energy transfer (FRET) pair whose fluorescent signal reports on the dimensions of the LEA sequence. We measure ensemble sensitivity calculated as the difference in FRET efficiency (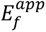) between a construct in buffer and under various solution conditions (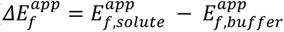). To contextualize these changes, we compared them to a homopolymer of equivalent length composed of glycine-serine (GS) repeats placed between the same FRET pair (Moses et al., 2020). Previous studies have shown that GS repeats behave as ideal homopolymers without structural biases, making them a good reference for SSS (Moses et al., 2024; Sørensen & Kjaergaard, 2019).

We cloned the open reading frames encoding the selected LEA proteins and fused them individually between mTurquoise2 and mNeonGreen fluorescent proteins. The recombinant proteins were expressed, purified (**Figure S2**) and analyzed using FRET to assess their conformational ensembles. We measured FRET efficiencies in neat buffer (20 mM sodium phosphate, 100 mM NaCl, pH 7.4) and in the same buffer supplemented with increasing concentrations of polyols (ethylene glycol, glycerol, sorbitol), sugars (sucrose, trehalose), amines (glycine, sarcosine, betaine), salts (NaCl, KCl), macromolecular crowders including Ficoll and polyethylene glycol (PEG) of different sizes (PEG200, PEG400, PEG1500, PEG2000, PEG4000, PEG6000, PEG8000, PEG10000), and denaturants (urea, guanidinium chloride).

We obtained 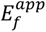, which showed that in buffer, AtEM1, AtLEA7, and AtLEA4-5 formed ensembles that were either more compact or similar in size to their corresponding GS repeats (**Figure 4a**). In contrast, the AtLEA6-2.2 ensemble was more expanded than its equivalent GS repeat (**Figure 4a**). This difference in ensemble dimensions may be attributed to the higher content of proline and/or aspartic acid residues (helix breaking residues) in AtLEA6-2.2 relative to the other tested proteins, consistent with the AlphaFold2 predictions (**Figure 1d**, **Figure 4b**).

**FIGURE 4.**
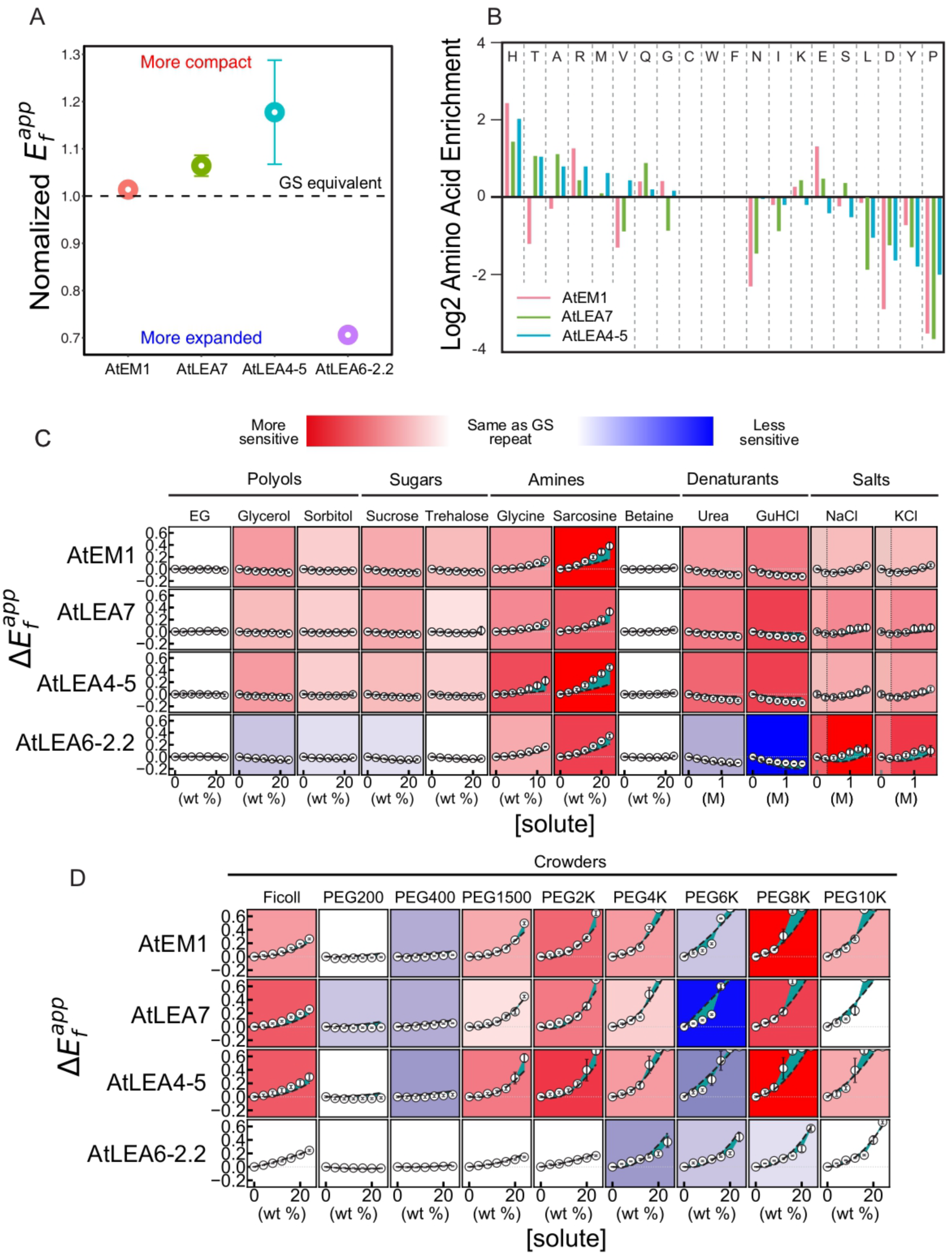
The conformational ensembles of Arabidopsis LEA proteins are sensitive to changes in solution chemistry *in vitro*. A) Average 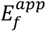 of AtEM1, AtLEA7, AtLEA4-5, and AtLEA6-2.2 in buffer solution normalized to a GS-repeat sequence of the same length (black dashed line). Markers show the mean and error bars are the SD from *n* = 3 repeats. B) Amino acid enrichment of AtEM1, AtLEA7, and AtLEA4-5 relative to AtLEA6-2.2. Positive values indicate that the composition of the amino acid is higher in the corresponding LEA protein than in AtLEA6-2.2. Negative values indicate that the composition of the amino acid is higher in AtLEA6-2.2 than in the corresponding LEA protein. C) Structural ensemble sensitivity (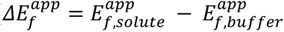) of AtEM1, AtLEA7, AtLEA4-5, and AtLEA6-2.2 in response to polyols, sugars, amines, denaturants, and salts. Black dashed lines are interpolated 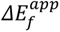 of GS-repeat sequences of equivalent lengths of each LEA protein. Cyan shaded regions are differences between 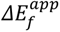 of LEAs and GS-repeat sequences. Background color of each plot indicates the structural sensitivity to that solute, with red/white/blue representing more/same/less sensitivity relative to the corresponding GS-repeat sequence, respectively. Data points are the mean and error bars are SD from *n* = 3 repeats. D) Same as (C) in response to macromolecular crowders. Data points are the mean and error bars are SD from *n* = 3 repeats.

Next, we added different osmolytes and calculated the change in apparent FRET efficiency (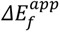) for each LEA protein. This allowed us to determine whether a sequence was more (red background) or less structurally sensitive (blue background) to a given osmolyte, and whether that osmolyte promoted compaction (positive 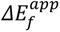) or expansion (negative 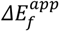) of the ensemble. We found that ensemble sensitivity varied relative to GS-repeat sequences of equivalent lengths, depending on both the identity of the LEA protein and the osmolyte added to the solution (**Figure 4c-d**). AtEM1, AtLEA7, and AtLEA4-5 were highly sensitive, despite having compact ensembles in buffer (**Figure 4a,c-d**), in agreement with our *in silico* observations (**Figure 3**). These ensembles were sensitive to amines, denaturants, and high molecular weight polymeric crowders (**Figure 4c-d**). AtLEA6-2.2 had the least sensitive ensemble of all, being insensitive (and in some cases less sensitive than equivalent GS-repeats) to most of the solutes except for amines and salts (**Figure 4c-d**). Overall, the ensembles of the four Arabidopsis LEA proteins examined displayed different levels of structural sensitivity to changes in the solution environment *in vitro*.

### 2.4 Arabidopsis LEA proteins exhibit unique patterns of chemical-specific ensemble sensitivity in solution

We next performed a comparative analysis of the effects of the different solute classes on LEA ensemble dimensions. To compare sensitivity differences among all LEA proteins in response to a particular osmolyte, we calculated 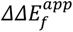 at the highest concentration of each osmolyte. 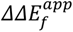 represents the difference in structural sensitivity (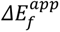) between each LEA protein and a GS repeat with the same number of amino acids (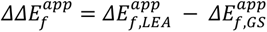). The first solute class we examined, amines, consists of organic molecules that can function as osmolytes and may be taken up by the cell upon hyperosmotic stress (Kaur et al., 2024). All tested amines, particularly glycine and sarcosine, induced compaction of LEA proteins’ ensembles (**Figure 4c**). However, we did not observe significant differences in structural sensitivity to amines among the different LEA proteins (**Figure 5a-c**).

**FIGURE 5.**
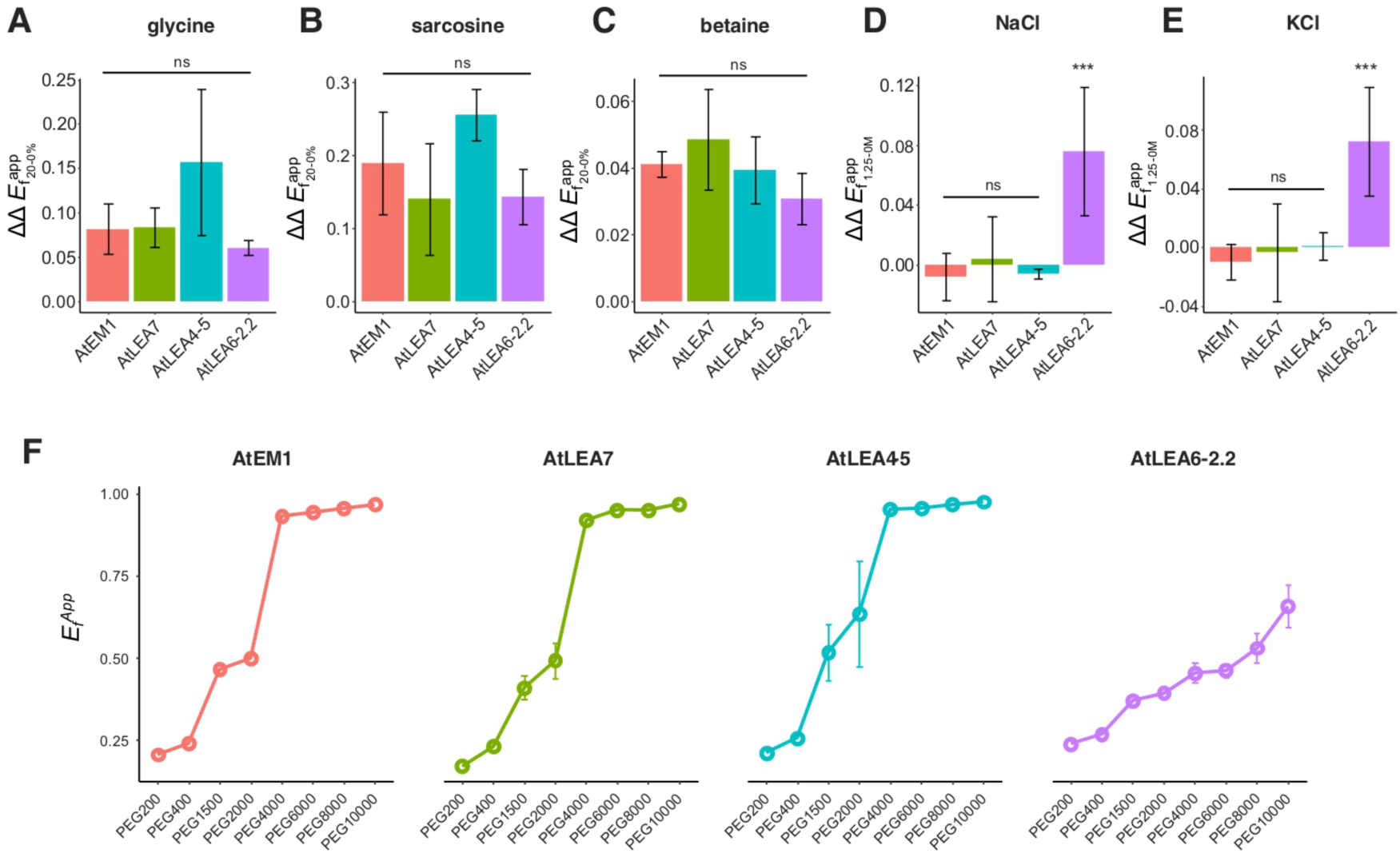
Arabidopsis LEA proteins exhibit unique patterns of chemical-specific ensemble sensitivity in solution. The difference in structural sensitivity compared to a GS repeat (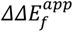) of AtEM1, AtLEA7, AtLEA4-5, and AtLEA6-2.2 in A) glycine (20-0%), B) sarcosine (20-0%), C) betaine (20-0%), D) NaCl (1.25-0M), and E) KCl (1.25-0M). Bars are the mean and error bars are SD from *n* = 3 independent measurements. One-way ANOVA with Tukey’s HSD test. \**p*<0.05, \*\**p*<0.01, \*\*\**p*<0.001, ns *p*>0.05. F) 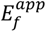 of AtEM1, AtLEA7, AtLEA4-5, and AtLEA6-2.2 after the addition of 20% PEG of the indicated size. Markers are mean and error bars are SD from *n* = 3 independent measurements.

The second solute class we looked at, salts, comprises some of the most relevant chemical species within a cell (Lang, 2007; Romero-Perez et al., 2023). Because salts can accumulate to even higher concentrations when cells undergo hyperosmotically-induced volume change, they play a significant role in modulating protein behavior (Liu et al., 2017). Their non-monotonic effect on LEA proteins (**Fig. 4c**) was reported for other disordered proteins as well (Vancraenenbroeck et al., 2019). This can be explained by the opposite effects of increasing electrostatic screening driving expansion at lower concentrations and increasing osmotic pressure which drives compaction at higher concentrations (Record et al., 1998). Strikingly, unlike all other osmolytes tested, the ensemble of AtLEA6-2.2 showed the highest structural sensitivity to high concentrations of NaCl or KCl (**Figure 5d-e**). This result highlights the divergent behavior that the ensembles of LEA proteins from different groups exhibit under specific chemical environments.

Cells are densely packed with macromolecules, and hyperosmotic stress or extreme desiccation massively increases macromolecular crowding (Meneses-Reyes et al., 2024). We found that most LEA proteins were highly sensitive to polymeric crowders *in vitro* (**Figure 4d**). To further characterize this effect, we compared 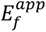 of each LEA as a function of the size of the PEG used to induce macromolecular crowding at a fixed concentration of the crowder (20% w/v). All LEA ensembles compacted in a PEG size-dependent manner, however, in contrast to the observation in salts, AtLEA6-2.2 was the least sensitive to this effect (**Figure 5f**).

Taken together, our *in silico* and *in vitro* approaches identified distinct signatures of structural sensitivity in LEA proteins in response to environmental perturbations. These findings suggest that specific attractive or repulsive interactions between the solution and the protein backbone modulate the effect of sequence-encoded conformational biases in this group of plant IDPs.

### 2.5 Arabidopsis LEA proteins gain helicity under desiccation

To investigate how the conformational changes observed with our FRET system in different solutions relate to residual structural elements, we evaluated the secondary structure of unlabeled (without fluorescent proteins) LEA proteins using circular dichroism (CD) spectroscopy. Consistent with predictions (**Figure 1d**), all LEA proteins tested were disordered in aqueous solution, as indicated by a pronounced minimum at around 200 nm in the CD spectra (**Figure 6**).

**FIGURE 6.**
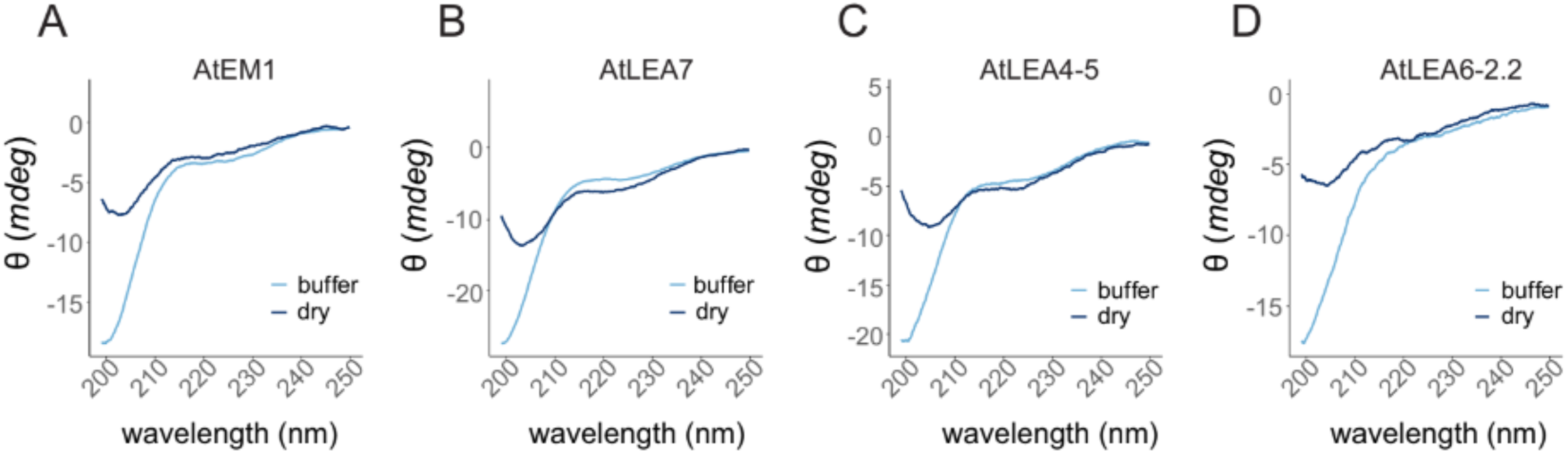
Arabidopsis LEA proteins gain helicity under desiccation. CD spectroscopy in buffer or in a desiccated state (dry) of A) AtEM1, B) AtLEA7, C) AtLEA4-5, or D) AtLEA6-2.2.

Next, we measured residual secondary structure under desiccation and performed Dichroweb deconvolution analysis with CDSSTR algorithm (Miles et al., 2022) to quantify the residual structural elements. Desiccation induced helicity in AtEM1 (14% gain), AtLEA7 (40% gain), and AtLEA4-5 (41% gain) (**Figure 6a-c, Figure S3**). Notably, the acquisition of helical conformations under desiccation was partially reversible, since rehydration of the samples resulted in more disorder (**Figure S3**). In contrast, AtLEA6-2.2 remained largely disordered with a minimal gain in helicity (7% gain) (**Figure 6d, Figure S3**).

These results suggest that sensitive LEA protein ensembles respond to desiccated conditions by gaining different levels of helicity, resulting in more compact ensembles under these conditions as compared to in buffer. Moreover, similar to its protective role, the sensing function of the LEA proteins tested appears to be controlled by the ensemble features encoded in their primary amino acid sequence.

## 3. DISCUSSION

The conformational ensembles of IDPs are highly sensitive to changes in solution properties, which has led to their proposed role as biological sensors of the cellular environment (Cuevas-Velazquez et al., 2021; Moses et al., 2023). In a previous study, we used the Arabidopsis disordered LEA protein AtLEA4-5 to develop a genetically-encoded fluorescent biosensor that reports cell volume changes during hyperosmotic stress in living cells across evolutionarily distant organisms. This demonstrated the high sensitivity of plant IDP ensembles, particularly LEA proteins, to environmental changes (Cuevas-Velazquez et al., 2021).

In this work, we investigated and compared the structural sensitivity of four Arabidopsis LEA proteins from different groups. We found that Arabidopsis LEA proteins from group 1 (AtEM1) and group 3 (AtLEA7) were as sensitive as AtLEA4-5 to changes in solution chemistry, both *in silico* and *in vitro*. Specifically, the ensembles of AtLEA4-5, AtEM1, and AtLEA7 compacted in response to increasing macromolecular crowding and increasing concentrations of amines. In contrast, the ensemble of AtLEA6-2.2, a group 6 LEA protein, was overall less sensitive or even insensitive to these conditions, but exhibited greater compaction in response to high salt concentrations. The contrasting sensitivities observed for these LEA protein families correlate with their ability to form helical conformations in the desiccated state. AtLEA4-5 was previously shown to form helices in high-osmolarity solutions (glycerol or ethylene glycol) (Cuevas-Velazquez et al., 2016; Rendón-Luna et al., 2024). This helical transition ability appears to be restricted to the N-terminal conserved region of group 4 LEA proteins from various plants. On the other hand, the group 6 LEA protein from common bean (PvLEA18) does not gain significant levels of helicity in high-osmolarity or macromolecular crowding solutions (Rivera-Najera et al., 2014), aligning with with our SSS results.

A closer look into the sequence-encoded properties of the LEA proteins we tested suggests that the sensitivity and the ability to gain helicity are determined by their primary amino acid sequence and composition. The high content of helix-breaking residues across the sequence of AtLEA6-2.2 correlates with its low helicity prediction and its more expanded ensemble in buffer conditions, relative to a GS-repeat sequence of equivalent length. Of particular interest is the greater sensitivity of AtLEA6-2.2 to salts, compared to the other LEAs. This finding suggests that short and/or long range ionic interactions might have a strong contribution to the ensemble properties of AtLEA6-2.2. Examining its charged residue composition, we found that aspartic acid (negatively charged) is more abundant in AtLEA6-2.2, whereas arginine (positively charged) is less abundant compared to the other three LEA proteins. This suggests that differences in the relative abundance of these amino acids contribute to AtLEA6-2.2’s salt sensitivity. Furthermore, this raises the possibility that pH may have a role in the protective and sensing functions of LEA proteins, as recently demonstrated for an Arabidopsis homologue of AtLEA6-2.2 during *in vitro* protection of LDH (Arroyo-Mosso et al., 2025).

Evidence suggests that LEA proteins from different groups can all function as biomolecular protectants, stabilizing enzymes and cellular structures under water limiting conditions (Cuevas-Velazquez et al., 2014; Hernández-Sánchez et al., 2024; Reyes et al., 2008). *In vitro*, many LEA proteins prevent the inactivation of labile enzymes such as LDH from the deleterious effects of desiccation. This protective function extends to IDPs found in other desiccation-tolerant organisms such as tardigrades (Boothby et al., 2017; Sanchez-Martinez et al., 2024). In this study, we showed that despite similarities in their amino acid composition, AtEM1, AtLEA7, AtLEA4-5, and AtLEA6-2.2 protect LDH from desiccation with slightly different efficiencies, suggesting that specific ensemble properties contribute to this function.

It has been shown that different factors contribute to the desiccation-protective function of IDPs *in vitro.* These include the ability to gain helicity, the self-assembly into homo- and hetero-oligomers, the presence and the number of specific sequence motifs (e.g., molecular recognition features, MoRFs), ensemble properties such as the radius of gyration, protectant-client interactions, and synergy with specific osmolytes (Biswas et al., 2024; Chakrabortee et al., 2012; Hernández-Sánchez et al., 2024; Kc et al., 2024; Nicholson et al., 2025; Rendón-Luna et al., 2024; Romero-Pérez et al., 2024; Sanchez-Martinez et al., 2024). We did not find a clear correlation between the protective efficiency and different ensemble properties of the LEA proteins tested here, including their structural sensitivity to the environment, so the extent to which ensemble properties encoded in the primary amino acid sequence influence the *in vitro* protective functions of different LEA proteins remains to be elucidated.

Overall, our results highlight the functional diversity of Arabidopsis LEA proteins across different groups and suggest that sequence encoded ensemble properties relate to these functions. While assessing functions of LEA proteins using *in vitro* approaches provide valuable insights, they also come with limitations, as discussed below. Our results show that structural sensitivity of desiccation-related IDPs to their environment correlates with their protective capacity for some, but not all LEA proteins. The extent to which ensemble sensitivity contributes to LEA protein function *in vivo* remains to be tested and will require the implementation of new methods to probe the function in a physiologically relevant context. Overall, identifying the molecular features encoded in the primary amino acid sequence that dictate sensitivity to specific physicochemical environments and/or desiccation protection, will not only aid in understanding how LEA proteins sense and respond to their surroundings, but would also help to design new proteins with improved sensing and/or protective capabilities, with potential applications in biotechnology.

### 3.1 Limitations and drawbacks of the study

A relevant aspect to consider for the interpretation of our results is that we used recombinant LEA proteins fused to a His-tag and a 12-residue scar at their N-terminal end. We acknowledge that the presence of a His-tag in our protective and secondary structure measurements might contribute to the observed results. Nevertheless, LEA proteins have been successfully characterized in their *in vitro* protective and structural properties using tagged versions (Artur et al., 2019; Goyal et al., 2005; LeBlanc & Hand, 2021; Navarro-Retamal et al., 2016; Sasaki et al., 2014). For example, when tagged and untagged versions of the LEA protein COR15A were compared using far-UV CD in hydrated and dry conditions, their ability to gain helicity was not significantly different (Navarro-Retamal et al., 2016). However, other studies have reported that the presence of a His-tag can affect the dynamics of IDPs, either by locally constraining the mobility of residues close to the tag or by exerting a more global impact (Mompeán et al., 2021; Walter et al., 2019). Additionally, the charge state of histidine residues could be modulated by pH changes during desiccation, which might have an impact in the protective function (pKa of lateral group = 6.0). In a recent work, Arroyo-Mosso and colleagues showed that AtLEA6-2.1, one of the Arabidopsis homologues of AtLEA6-2.2, efficiently protects LDH from desiccation (10:1 LEA to LDH molar ratio) at pH 8, but not at pH 7.5 or 6.8, suggesting that the physicochemical properties of the surrounding environment and their effect on certain amino acid residues contribute to the protective function of LEA proteins. Overall, the presence of a His-tag might not robustly alter how LEA proteins behave in solution, as revealed by the differential behaviours observed in our study; however, appropriate caution should be exercised.

## 4. MATERIALS AND METHODS

### 4.1 Sequence analysis

Hydropathy, FCR and NCPR values were calculated with localCIDER v0.1.13. Disorder and three-dimensional structure prediction were generated with metapredict online v2.0 and AlphaFold2 (ColabFold v1.5.5), respectively. ALBATROSS-Colab v1.0.3 was used for prediction of ensemble dimensions.

### 4.2 Transcriptomic analysis

Raw RNA-seq data were cleaned using fastp v0.20.0, removing reads shorter than 33 bp and those with a quality score below Q20 across 40% or more of their sequence. Bowtie2 v2.4.4 was used for local alignment to the reference Arabidopsis genome (GenBank: GCA_000001735.2), retaining uniquely mapped reads. Annotation and read counting were performed using featureCounts v2.0.0 with default options. Differential expression analysis of Arabidopsis seeds was performed using DESeq2 (Galaxy v2.11.40.6+galaxy2) with default parameters, except for the outlier filtering step. Raw RNA-seq data from seeds before (silique) and after (dry seeds) desiccation, was obtained from BioProject PRJNA314076.

### 4.3 All-atom simulations

All-atom Monte Carlo simulations of LEA proteins were conducted using the Solution Space Scanning method described in previous studies (Moses et al., 2020, 2024). These simulations employed a modified ABSINTH implicit solvent force field with the CAMPARI v2_09052017 simulation suite, which has been extensively benchmarked against experimental data.

Briefly, the overall Hamiltonian can be written as follows:

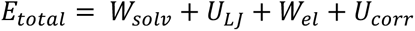

Here, *W_solv_* is the energy term describing the interaction between the protein surface and the surrounding solution environment. Seven different solution conditions were simulated by varying the *W_solv_* by up to 6% (+3% to −3% relative to the buffer condition). This approach enabled the force field to capture changes in the repulsive interactions between the solution and proteins triggered by desiccation.

For each solution condition, five independent conformational trajectories were generated for each protein, following 1×10^7^ steps of equilibration at 310 K. Each trajectory contained 5,600 conformations in total.

The normalized radius of gyration and end-to-end distance were calculated with custom Python scripts based on the MDtraj library. Normalization was performed relative to the LEA protein ensemble under the most expanded solution environment.

### 4.4 Construct design and cloning

The open reading frames (ORFs) that code for the selected LEA proteins (AtEM1: At3g51810; AtLEA7: At1g52690; AtLEA4-5: At5g06760; AtLEA6-2.2: At2g23120) were synthesized by GenScript and inserted into the pET28a plasmid, widely used for the expression of recombinant proteins in *Escherichia coli* (*E. coli*). This plasmid includes the inducible T7 promoter, an N-terminal 6xHis affinity tag for protein purification by affinity chromatography, a kanamycin resistance gene, and the genes encoding the LEA proteins. For FRET experiments, the genes encoding the LEA proteins are located between genes encoding the fluorescent proteins (mTurquoise2 and mNeonGreen). For expression of unlabelled proteins, the open reading frames of the LEA proteins were synthesized by GenScript and inserted into the same plasmid, excluding fluorescent proteins.

### 4.5 Expression and Purification of unlabeled recombinant proteins

The plasmids encoding for each of the LEA proteins were transformed in BL21 (DE3) *E. coli* (New England Biolabs) and plated on LB agar plates containing 50 µg/mL kanamycin. Cultures were incubated in LB medium with 50 µg/mL kanamycin at 37°C while shaking at 180 rpm until an optical density at 600 nm (OD600) of 0.6 was reached. Protein expression was induced with 1 mM isopropyl-D-1-thiogalactopyranoside (IPTG). 4 hours after the induction, cells were harvested by centrifugation at 4,000 rpm for 30 minutes at 4 °C. Cell pellets were resuspended using 5 mL of lysis buffer (50 mM NaH_2_PO_4_, 300 mM NaCl, 10 mM imidazole, pH 8.0). Resuspended samples were stored at −80 °C. For purification, samples were thawed at room temperature and heat lysis was performed for 15 min using boiling water. Insoluble debris were removed by centrifugation at 10,500 rpm at 10 °C for 30 min. Supernatant was loaded onto a column packed with Ni-NTA beads (G-Biosciences) for purification. LEA protein constructs were eluted using an elution buffer (50 mM NaH_2_PO_4_, 300 mM NaCl, 250 mM imidazole, pH 8.0). Constructs were concentrated using an Amicon Ultra-15 (3 kDa) tube and desalted using disposable PD-10 desalting columns (cytiva). Purified LEA proteins were then quantified fluorometrically (Qubit4 Fluorometer, Invitrogen), flash frozen and lyophilized (FreeZone 6, Labconco) for 48 h. Lyophilized LEA proteins were stored at −20 °C.

Final storage conditions varied according to the assay requirements. For SSS, desalted proteins were stored in 50 mM NaH_2_PO_4_ buffer (pH 8.0) at 4°C for immediate use. For the LDH protection assay, purified proteins were flash-frozen in liquid nitrogen, lyophilized for 48 hours using a laboratory freeze dryer, and stored at −20°C until further analysis.

### 4.6 Lactate Dehydrogenase (LDH) desiccation protection assay

Assays were performed in triplicate following published protocols (Boothby et al., 2017). LEA proteins were resuspended in a concentration range from 0.01 mg/mL to 10 mg/mL with resuspension buffer (25 mM Tris, pH 7.0). L-LDH from rabbit muscle (Sigma-Aldrich Cat #10127230001) was added to the sample at 0.1 g/L (2.86 µM considering a monomer).The final volume of each sample was 100 µL. Half of the sample was stored at 4 °C while the other half was desiccated for 16 h (SAVANT Speed Vac Concentrator). After 16 hours, hydrated and desiccated samples were brought to a volume of 250 µL with water. In a quartz cuvette, a mixture of 10 µL sample, 980 µL phosphate buffer (100 mM Sodium Phosphate, 2 mM Sodium Pyruvate [Sigma-Aldrich]) and 10 µL of 10 mM NADH [Sigma-Aldrich], pH 6 was made to measure LDH activity. Measurement of the conversion of NADH to NAD+ was conducted by absorbance reading at 340 nm for 1 minute using the UV-Vis function of NanodropOne (Thermo Scientific). The percent protection of LDH activity comes from the quotient of the initial, linear reaction rate of each desiccated sample divided by its corresponding hydrated control. Each sample was performed in triplicate.

### 4.7 Expression and purification of recombinant proteins fused to fluorescent proteins

The plasmids encoding for each of the LEA proteins were transformed into the BL21 (DE3) pLysS competent cells (Promega) strain of *E. coli* using the heat shock transformation method. Cultures were incubated in LB medium with kanamycin (50 μg/mL) at 37°C with shaking at 225 rpm until reaching an optical density at 600 nm (OD600) of 0.6. Protein expression was induced by adding 1 mM IPTG. Cells were harvested by centrifugation for 15 minutes at 3,000 rcf, supernatant was discarded, and cell lysis was performed using a QSonica Q700 Sonicator (QSonica, Newtown, CT) and lysis buffer (50 mM NaH_2_PO_4_, 0.5 M NaCl, pH 8.0). After a second centrifugation of the cell lysate for 1 hour at 20,000 rcf, the supernatant was loaded onto a column packed with Ni-NTA beads (Qiagen) for purification. Elution of each construct was carried out using an elution buffer (50 mM NaH_2_PO_4_, 0.5 M NaCl, 250 mM imidazole, pH 8.0). Constructs were concentrated using an Amicon Ultra-15 (10 kDa) tube and centrifuged for 7 minutes at 4°C. Subsequently, the concentrated constructs were purified by high-performance liquid chromatography (HPLC) using a HiLoadTM 16/600 SuperdexTM 200 pg column with the Superdex 200 Hi Load 16 60 RUN program for 4 hours at 0.5 ml / min, to a total of 120 mL at −4°C. Finally, the purified constructs were divided into 200 μL aliquots, rapidly frozen in liquid nitrogen, and stored at −80°C. The concentration of purified LEA proteins was determined by UV-visible absorbance at 506 nm (peak absorbance wavelength for mNeonGreen), and purity was assessed by SDS-PAGE after thawing and before use.

### 4.8 Preparation of solutions for SSS

The solutions were prepared by dissolving each solute in buffer (20 mM NaH_2_PO_4_, 100 mM NaCl, pH 7.4). No additional salt was added to the NaCl and KCl solutions. The same buffer was used for all solutions. The solutes used were acquired from: Alfa Aesar (Sarcosine, PEG200, PEG400, PEG1500, PEG2000, PEG4000, PEG6000, PEG8000, PEG10000), VWR (D-sorbitol), GE Healthcare (Ficoll), TCI (D-(+)-trehalose dihydrate), Thermo Scientific (guanidine hydrochloride), Acros Organics (D-mannitol, betaine monohydrate), Sigma-Aldrich, and Fisher BioReagents (ethylene glycol, glycerol, glycine, L-proline, potassium chloride, sodium chloride, urea).

### 4.9 Solution space scanning experiments with FRET

Different amounts of 20 mM NaH_2_PO_4_ stock buffer, 100 mM NaCl, pH 7.4, and purified protein were placed in each well of black plastic 96-well plates (Nunc) to obtain a final volume of 150 μL. In each well, a final concentration of 1 μM protein and different concentrations of each solute were added. Fluorescence measurements were obtained using a CLARIOstar plate reader (BMG LABTECH), taken from above at a focal height of 5.7 mm, with a fixed gain of 1020. Three replicates were performed for each FRET construct in the stock buffer and in all solution conditions. Fluorescence emission spectra were obtained for each FRET construct by exciting the sample in a 16 nm band centered at 420 nm, with a dichroic at 436.5 nm, and measuring fluorescence emission from 450 to 600 nm, averaging over a 10 nm window moved at intervals of 0.5 nm. Donor and acceptor base spectra for each solution condition were obtained using the same excitation and emission parameters in solutions containing 1 μM of mTurquoise2 or mNeonGreen alone, and measuring fluorescence emission from 450 to 600 nm.

### 4.10 Calculation of FRET efficiencies

The apparent FRET efficiency (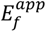) for each LEA protein construct in each condition was calculated by linear regression of the fluorescence spectrum of the construct with the separate emission spectra of the donor and acceptor in each condition (to correct for solute-dependent effects on the emission of the fluorescent proteins used). 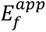 was calculated using the following equation:

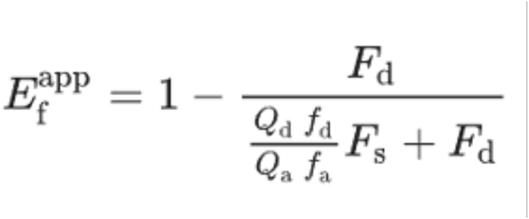

where *F*d is the decoupled donor contribution, *F*s is the decoupled acceptor contribution, *f*d is the area-normalized donor spectrum, *f*a is the area-normalized acceptor spectrum, *Q*d of 0.93 is the quantum yield of mTurquoise2 and *Q*a of 0.8 is the quantum yield of mNeonGreen. The data for each series of solution conditions consisting of increasing concentrations of a single solute were processed as described in (Moses et al., 2024).

### 4.11 Dry circular dichroism

Lyophilized recombinant proteins were weighed and resuspended in 25 mM Tris buffer, pH 7, to an approximate concentration of 100 μM. The concentration of the solution was confirmed using a Qubit (Life Technologies, Qubit 3.0 Fluorometer). A 20 μL aliquot of the protein sample was measured in the far UV range using a CD spectropolarimeter (JASCO J-1500 model) in a 0.05 mm quartz cuvette. A new aliquot was then deposited on one side of a quartz cuvette and spread across the sample window with a micropipette tip to an area of approximately 1 cm^2^. The sample was then desiccated in a vacuum chamber for one hour. Another CD measurement was taken as soon as the vacuum stopped. 20 μL of distilled water (MilliQ) was dispensed onto the dry sample, and the sample was left to rehydrate at room temperature for 5 minutes before a final measurement was taken. Each measurement was performed in triplicate.

### 4.12 Data visualization and statistical analysis

All experimental data were plotted using R-Studio v1.3.1073, Python v3.8.8 and GraphPad Prism v10.1.0. Differential expression analysis hypothesis testing was made using DEseq2 (Galaxy v2.11.40.6+galaxy2) default methods. Solution space scanning data were fitted to linear and quadratic models to explain variations in apparent FRET efficiency (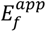) as a function of solute concentrations, utilizing the *curve_fit* function from SciPy. Statistical differences in LEA proteins’ 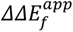 at the highest concentration of each osmolyte were analyzed using one-way ANOVA with Tukey’s post hoc test. LDH protection data was fitted into a sigmoidal curve by fitting a 5PL regression analysis using GraphPad Prism v10.1.0 from which the resulting PD50 values were derived. The statistical significance of LEA proteins’ PD50 values was assessed using one-way ANOVA followed by Tukey’s post hoc test. Circular dichroism data shown in the supplementary figures was deconvoluted and statistically analysed using Dichroweb server and independent two-sample t-test, respectively.

## ACKNOWLEDGEMENTS

This material is based upon work supported by a grant from the University of California Institute for Mexico and the USA (UC MEXUS) and the Consejo Nacional de Humanidades, Ciencias y Tecnología de México (CONAHCYT), project CN-20-113. We acknowledge the support from the Global Perspectives Grant program from the College of Agriculture, Life Sciences, and Natural Resources of the University of Wyoming. Support for this project came from the Consejo Nacional de Humanidades, Ciencias y Tecnología (CONAHCYT), project 252952; the Programa de Apoyo a Proyectos de Investigación e Innovación Tecnológica, Dirección General de Asuntos del Personal Académico, Universidad Nacional Autónoma de México (UNAM-PAPIIT), project IA203422; and Programa de Apoyo a la Investigación y el Posgrado, Facultad de Química, Universidad Nacional Autónoma de México, grant 5000-9182. L.D.P.-N. (CVU 1099541) and D.P.-V. (CVU 1224622) acknowledge CONAHCYT for their MSc fellowships. S.S, F.Y., and E.G. are funded by NIH grant R35GM137926. S.S. acknowledges support from the Alfred P. Sloan Foundation. S.S. and T.C.B. are part of the Water and Life Interface Institute (WALII), supported by NSF grant 2213983, and from NSF IntBIO grants 2128067 (SS) and 2128069 (TCB).

