## Supplemental Table 1 for "Functional diversity of Arabidopsis late embryogenesis abundant proteins in response to changes in the physicochemical environment"

**Table S1. Sequence-encoded properties of AtEM1, AtLEA7, AtLEA4-5, and AtLEA6-2.2**. FCR: Fraction of charged residues. NCPR: Net charge per residue. $\kappa$: kappa. All properties were calculated with CIDER.

| AGI code | Name | Disorder promoting | FCR | NCPR | $\kappa$ | hydropathy |
|---|---|---|---|---|---|---|
| At3g51810 | AtEM1 | 0.83 | 0.36 | -0.04 | 0.12 | 3.03 |
| At1g52690 | AtLEA7 | 0.88 | 0.29 | 0 | 0.06 | 3.18 |
| At5g06760 | AtLEA4-5 | 0.81 | 0.20 | 0.02 | 0.06 | 3.68 |
| At2g23120 | AtLEA6-2.2 | 0.86 | 0.24 | -0.07 | 0.11 | 3.49 |
