## Supplementary Information for "Functional diversity of Arabidopsis late embryogenesis abundant proteins in response to changes in the physicochemical environment"

### SUPPLEMENTARY MATERIAL

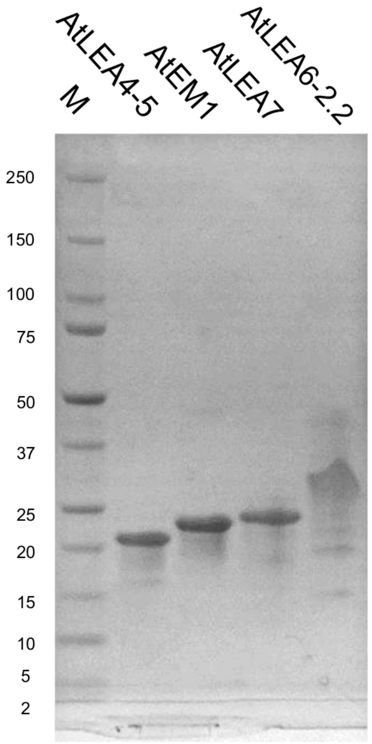


**FIGURE S1**. **Purified recombinant LEA proteins used in this study**. SDS-PAGE 4-20%. M: Molecular weight marker. We observed the abnormal mobility of purified proteins in SDS-PAGE as reported in other studies of LEA proteins and IDPs (Iakoucheva et al., 2001; Olvera-Carrillo et al., 2010; Rivera-Najera et al., 2014).

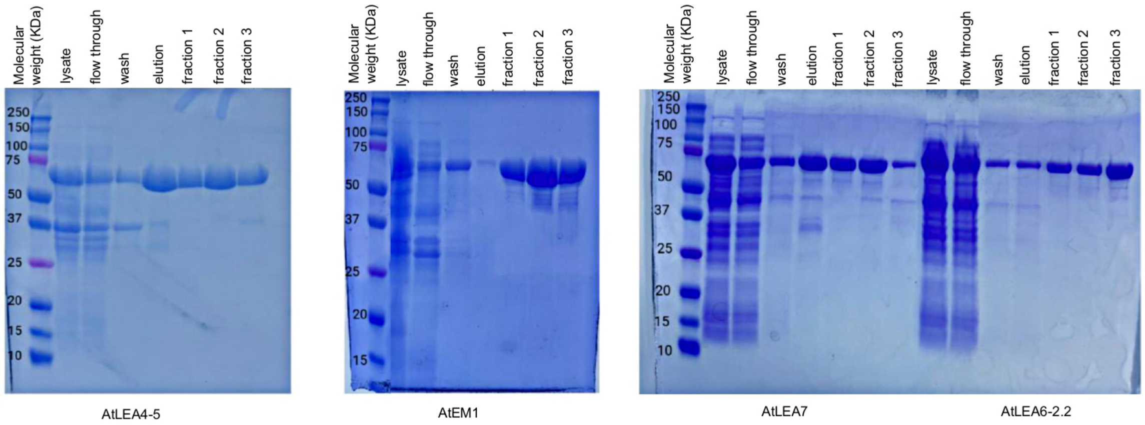


**FIGURE S2. Purification of recombinant LEA proteins fused to mTurquoise2 and mNeonGreen fluorescent proteins.** SDS-PAGE 4-20%. Fractions 1, 2, and 3 were combined for SSS measurements.

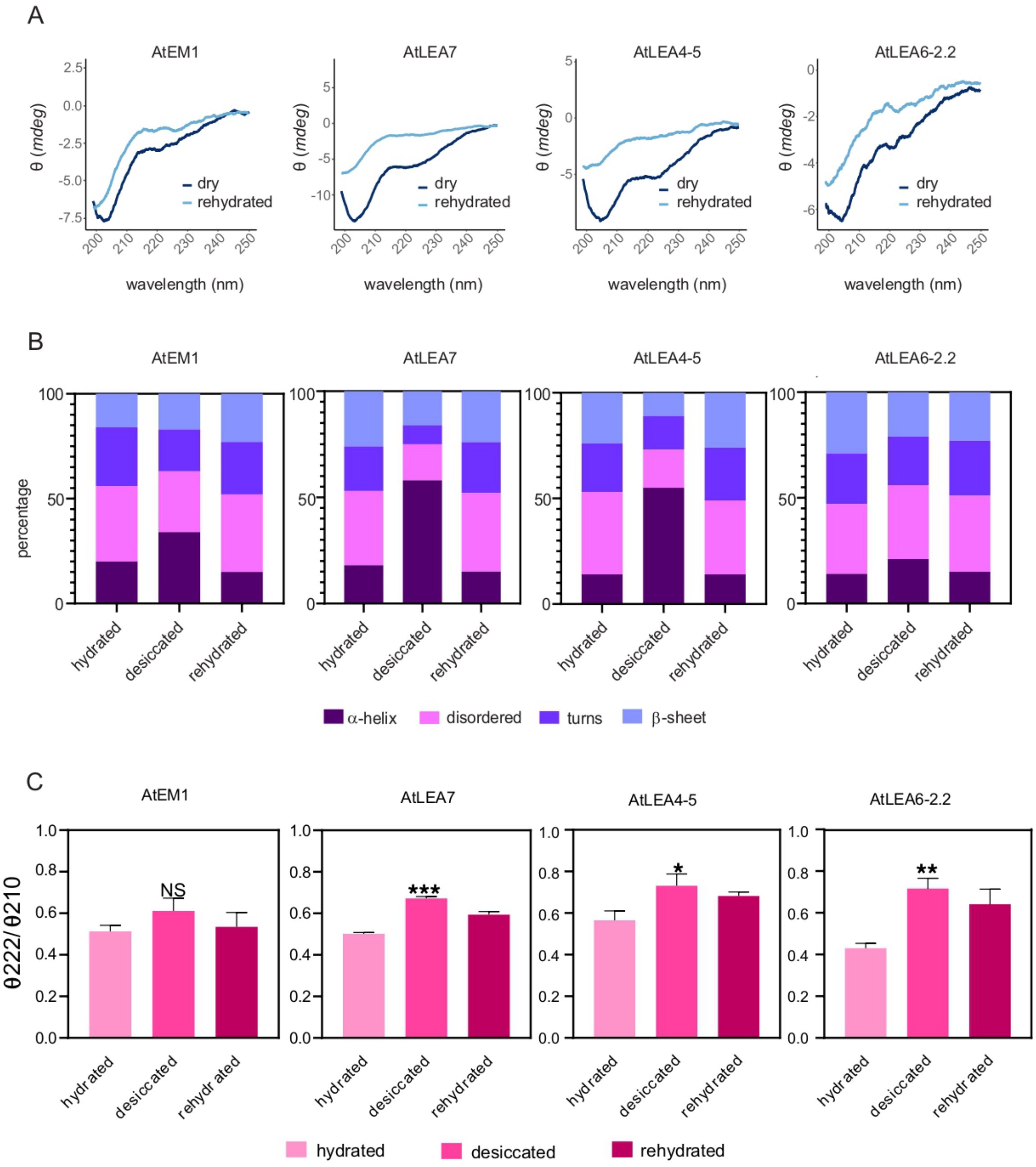


**FIGURE S3**. **Secondary structure analysis from CD measurements in desiccated samples**. A) Far UV-CD spectra of AtEM1, AtLEA7 AtLEA4-5, and AtLEA6-2.2 desiccated (dry, dark blue) and after rehydration of the sample (light blue). B) Secondary structure Dichroweb deconvolution of AtEM1, AtLEA7 AtLEA4-5, and AtLEA6-2.2 in buffer (hydrated), desiccated, and after rehydration of the sample. C) Ellipticity ratio 222 nm/210 nm of AtEM1, AtLEA7 AtLEA4-5, and AtLEA6-2.2 in buffer (hydrated), desiccated, and after rehydration of the sample. Independent two-sample t-test: *$p < 0.05$, **$p < 0.01$, ***$p < 0.001$, *ns* $p > 0.05$.
